# Local Haplotype Classifiers enable Efficient, Flexible, and Secure Genotype Imputation and Downstream Analyses

**DOI:** 10.1101/2024.12.01.626205

**Authors:** Muhammad Nadeem Cheema, Anam Nazir, Jungho Moon, Yongwoo Oh, Ardalan Naseri, Degui Zhi, Xiaoqian Jiang, Miran Kim, Arif Harmanci

## Abstract

The decreasing cost of genotyping technologies led to abundant availability and usage of genetic data. Although it offers many potentials for improving health and curing diseases, genetic data is highly intrusive in many aspects of individual privacy. Secure genotype analysis methods have been developed to perform numerous tasks such as genome-wide association studies, meta-analysis, kinship inference, and genotype imputation outsourcing. Here we present a new approach for using lightweight haplotype classifier models to use predicted haplotype information in a flexible privacy-preserving framework to perform genotype imputation and downstream tasks. Compared to the previous secure methods that rely main on linear models, our approach utilizes efficient models that rely on utilizing haplotypic information, which improves accuracy and increases the throughput of imputation by performing multiple imputations per model evaluation.

## Introduction

Genotype imputation[1–4] is used as an important intermediate step in genome-wide association testing, genetic risk score estimation, and ancestry analysis. The standard imputation methods rely on using diverse and large phased genotype panels as reference panels, e.g., The 1000 Genomes Project[5–7], Haplotype Reference Consortium[8], NHLBI Trans-Omics for Precision Medicine (TOPMed) panel[9]. Given an input genome that is sparsely genotyped (or typed) using, for example, genotyping arrays, the imputation tools compare the typed variant genotypes with the reference panel and aim to fill in the genotypes for the missing (untyped) variants. The state-of-the-art imputation methods first pre-phase the sparsely typed (which we refer to as input or query) panel to estimate a diploid representation of the input panel. Next, a hidden Markov model (formulated by Li-Stephens model[10]) is used to estimate the haplotype at which each typed variant resides on. These estimations are performed using highly optimized forward-backward algorithms[11–13], which is then used to assign the final allele probabilities and imputed dosages to all untyped variants. When the reference panel is representative of the haplotypic diversity of the input panel the imputation is highly accurate and can be used in downstream tasks. Thus, it is generally necessary to use large reference panels. Due to high computational requirements and protection of the reference panels, these are often outsourced to imputation servers[1,8,14,15]. The input panel, however, must still be submitted to the imputation server and may cause privacy concerns for the subjects in the query panel. We have organized one of the recent genomic privacy challenges to spearhead the development of secure genotype imputation services[16].

To protect the outsourcing, we and other groups have developed regression-based models and demonstrated that secure imputation outsourcing can be technically feasibly achieved using cryptographic approaches using Homomorphic Encryption (HE)[17–19] based primitives where the query data is encrypted in transit, at rest, and even while in analysis. In regression-based approaches, the idea is to impute each untyped variant using the typed variants in the vicinity of the untyped variant[20–22]. While these approaches work surprisingly well, the accuracy is limited for low-frequency and rare variants due to the simplicity of the linear models and the lack of training data for rare variants. Further, training becomes highly inefficient for large panels. More importantly, these imputation models do not explicitly make use of the haplotype information in the reference panel data.

Another approach proposed porting the imputation software source code into Intel’s Secure Guard eXtension (SGX)[23] enclaves and running the algorithms inside the enclaves. While this is an enticing option, SGX is far from becoming an industry standard, it has clear security flaws and is not adopted by all major cloud providers[24,25]. Since then, numerous regression-based methods have been developed with HE-based cryptographic protection schemes.

There are new efforts to build machine learning-based approaches that make use of deep neural networks to replace the reference panels with models. The query site can download the imputation model and perform imputation locally (model-to-data, M2D, philosophy[26]). This way no query data ever leaves the site, providing close-to-perfect privacy. While these methods are highly promising, they are strongly limited by hardware requirements with almost a unique reliance on very costly and specific GPU resources (similar to reliance on Intel’s SGX enclaves), especially in the training phase. Furthermore, these approaches require very large models to be transported and evaluated to the query site. This may not be always feasible due to network bandwidth requirements[27–29]. While deep learning approaches can protect query panels, recent studies indicate that the complex and large model may leak membership information[30], which is an unexplored risk that may affect model development efforts. In other words, the modes may leak information (via, for example, divergence attacks) about the subjects and their relatives in the reference panel, which leads to distinct privacy concerns. This may be a real issue for the deep learning models when model sizes surpass the size of the training data by, sometimes, a large margin[31].

In this study, we present a privacy-aware haplotype analysis framework that can be used for providing imputation services and downstream analysis tasks. To approach this problem, we break down the genotype imputation into two steps: The first step makes use of lightweight local haplotype classifier models, which assign the probabilities for haplotypes for the query subject. This is followed by linear projection of the haplotype predictions on the local haplotype matrix of the untyped variants to generate the final imputed genotypes. Compared to the per-untyped-variant regression-based approaches, this approach greatly decreases the number of models to be trained since number of local haplotype models we train is 2-3 orders of magnitude smaller than the total number of untyped variants in the reference panels. Compared to the deep learning-based *model-to-data* approaches, the local haplotype classifier models are again orders of magnitude smaller in size, they can be trained on CPUs without requiring substantial GPU resources and are much more feasible to be shared over network.

Our main objectives in this study are three-fold: We first present the local haplotype classifiers as a viable and flexible analysis framework that can enable efficient and flexible route for genomic analysis. We demonstrate that these models have much higher genotype imputation accuracy compared to the original linear models for imputation. Secondly, the flexibility of the presented approaches opens new interesting routes to protect data privacy at different levels of pipeline development and can alleviate regulatory requirements around data sharing. Motivated by the simplicity and flexibility of these models, we champion the development of “HE-convertible” models, i.e., the ease with which the haplotype classifier framework is compliant with HE-conversion of the models using HE-based primitives. We demonstrate that the haplotype classifier outputs can be used directly for downstream tasks such as collaborative genotype-phenotype analysis, e.g., association testing, and meta-analysis. Finally, we argue that our haplotype-based analysis framework for general genetic analysis with flexible routes for privacy protection and collaborative analysis in imputation and downstream analysis tasks.

## Results

We first describe the building and training of haplotype classifier models, we next assess the accuracy of imputation, and present downstream tasks.

### Local Haplotype Classifier Models

The input to model training is the phased reference panel (Fig. 1a). We first resample the phased reference panel to increase the haplotypic diversity of the panel using ProxyTyper[32]. The resampling is performed using the default sampling parameters of ProxyTyper.

**Fig 1.**
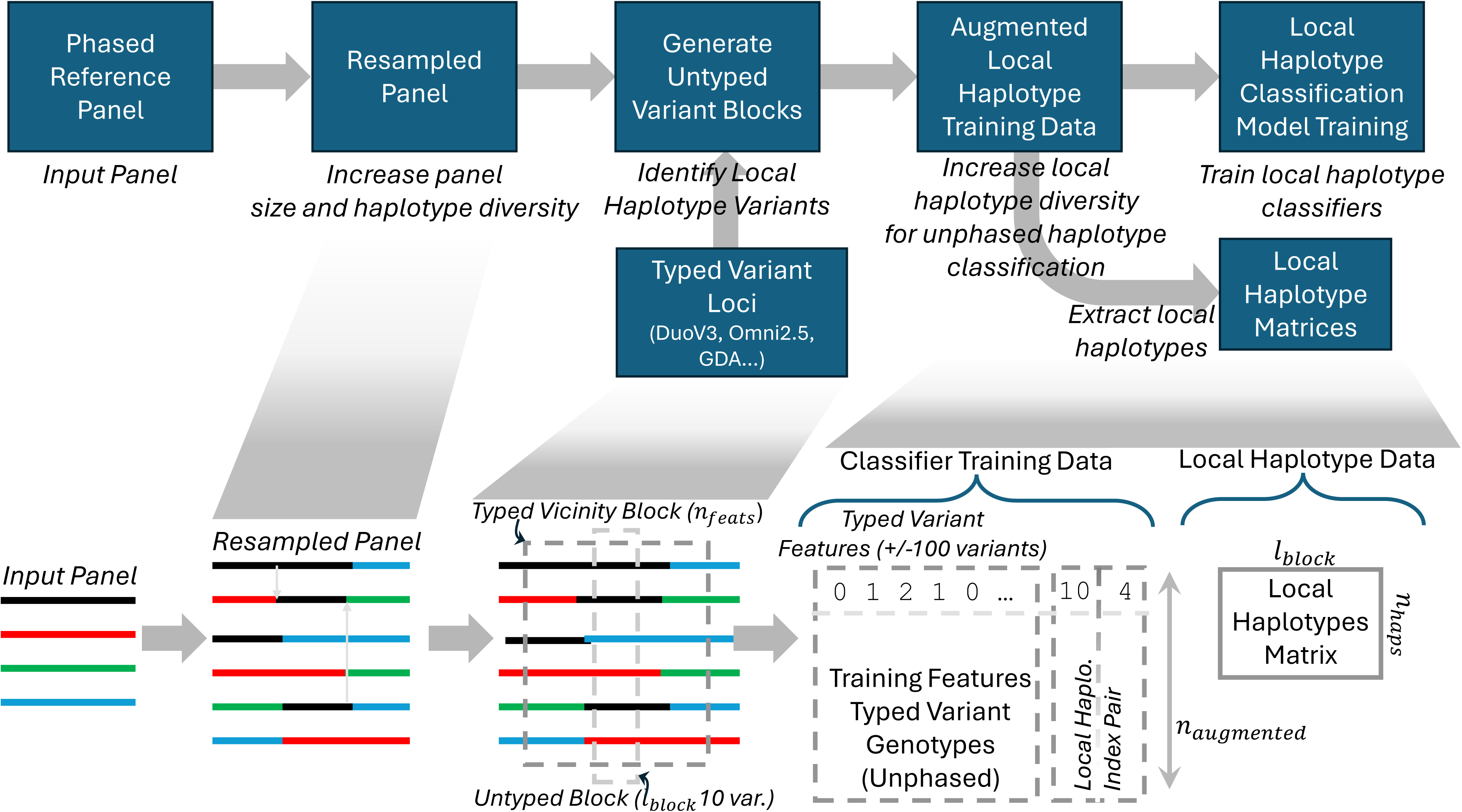
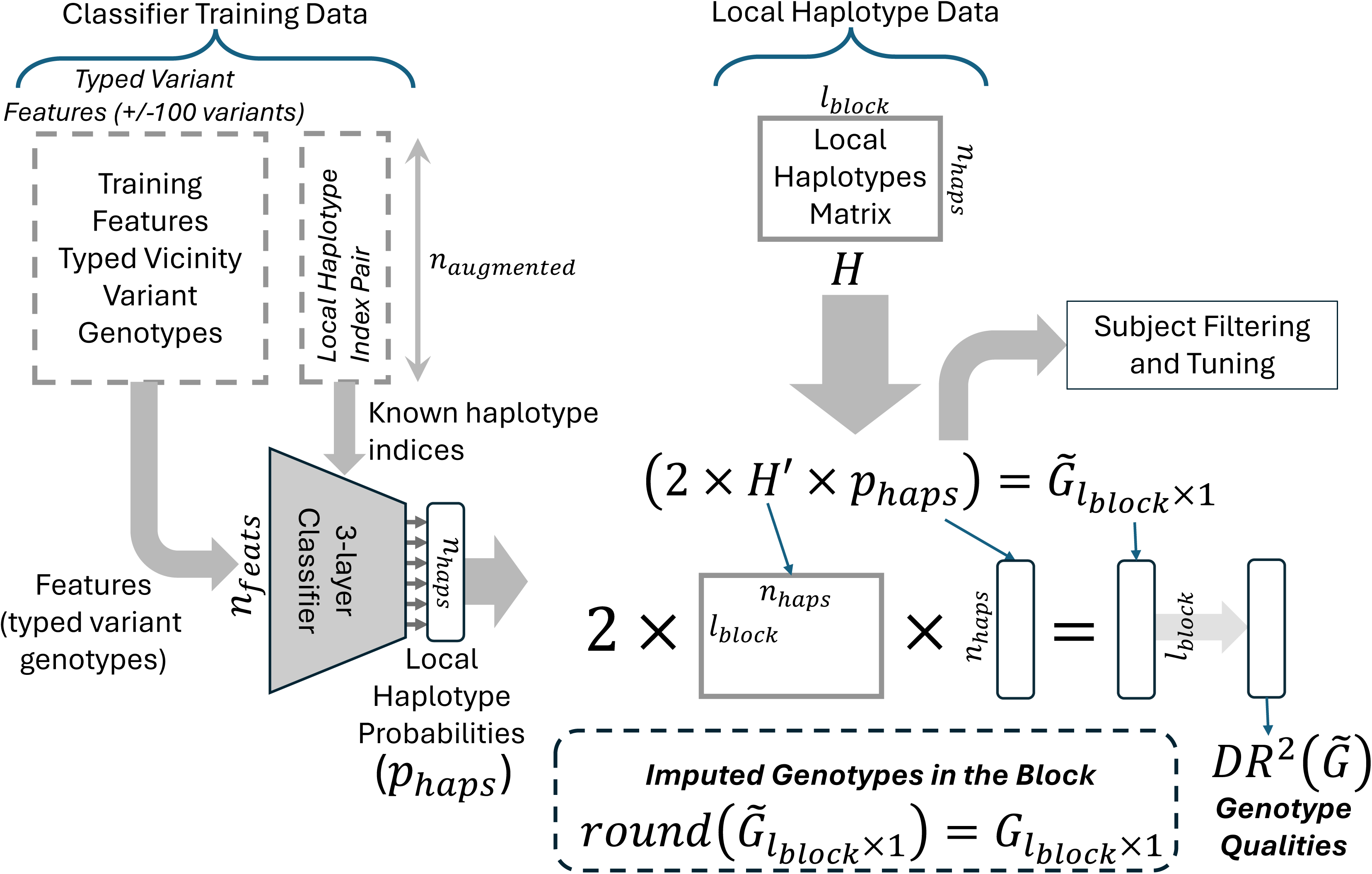
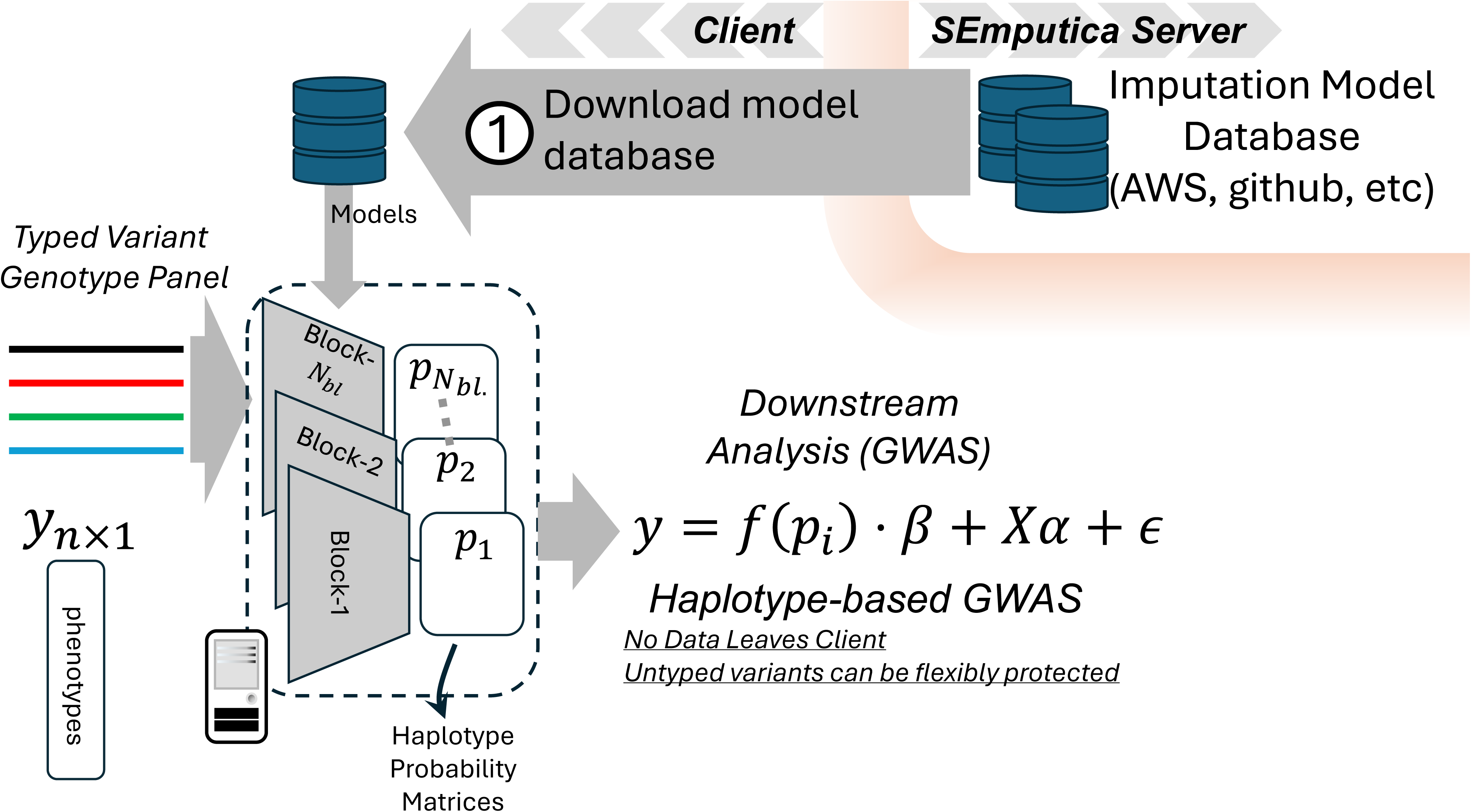
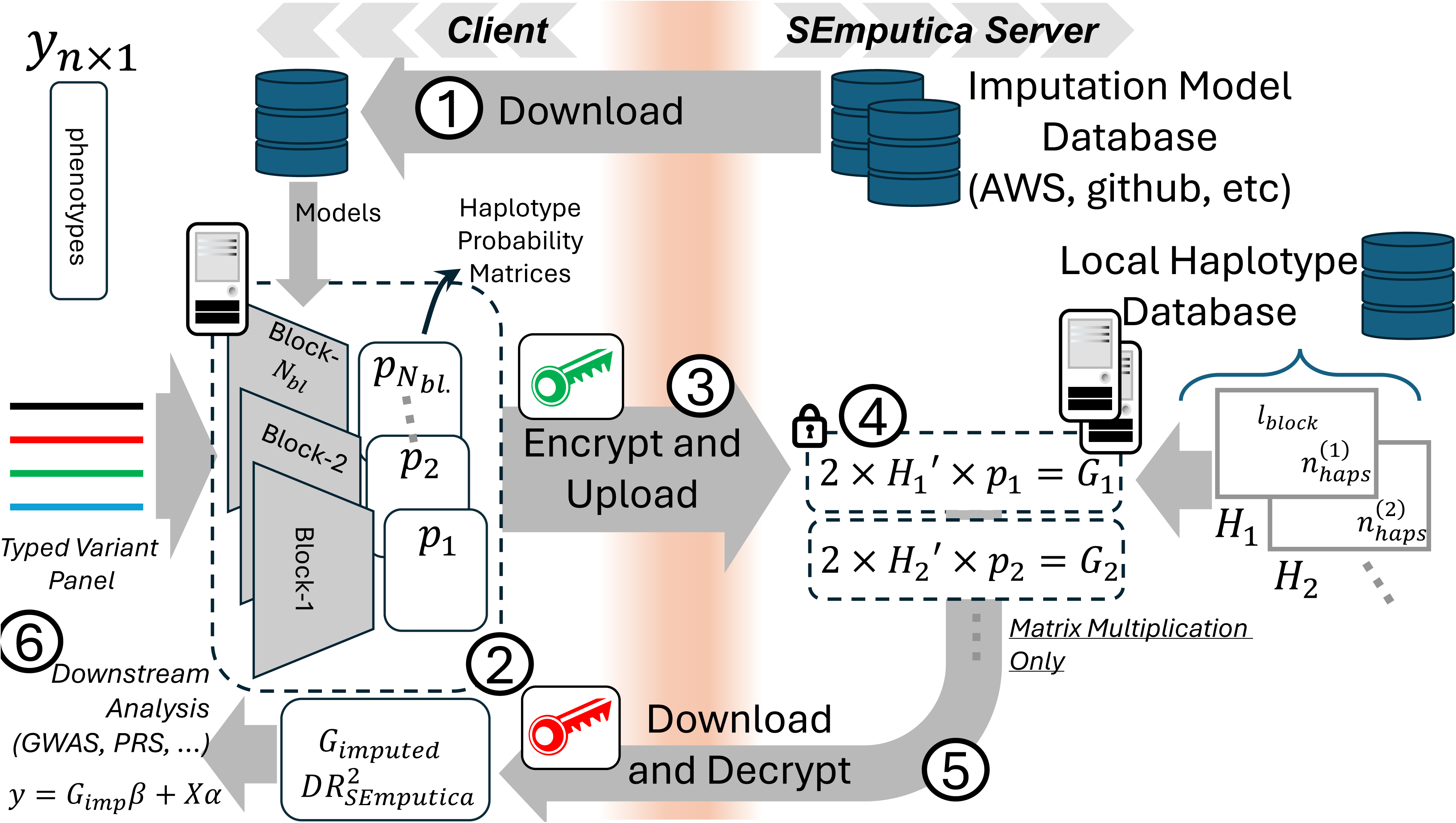

The typed variants are filtered with respect to reference minor allele frequency (MAF) cutoff, which is 1% by default. This operation removes some typed variants, which may decrease informative variants. We however observed that variants with low MAF tend not to provide informative features for the regression-based models.

Next, we use the untyped variants to define the local haplotype blocks on the genome. The untyped variants are first filtered with respect to reference MAF (1% by default). The genome is segmented into *n_block_*-long local haplotype blocks of the remaining untyped variants that we use to define local haplotypes. At each block, the local haplotypes are identified as the unique *n_block_* long allele sequences among the total set of allele sequences (Fig 2a). By default, we use *n_block_* is set to 10 (or maximum of 0.01 cM).

**Fig 2.**
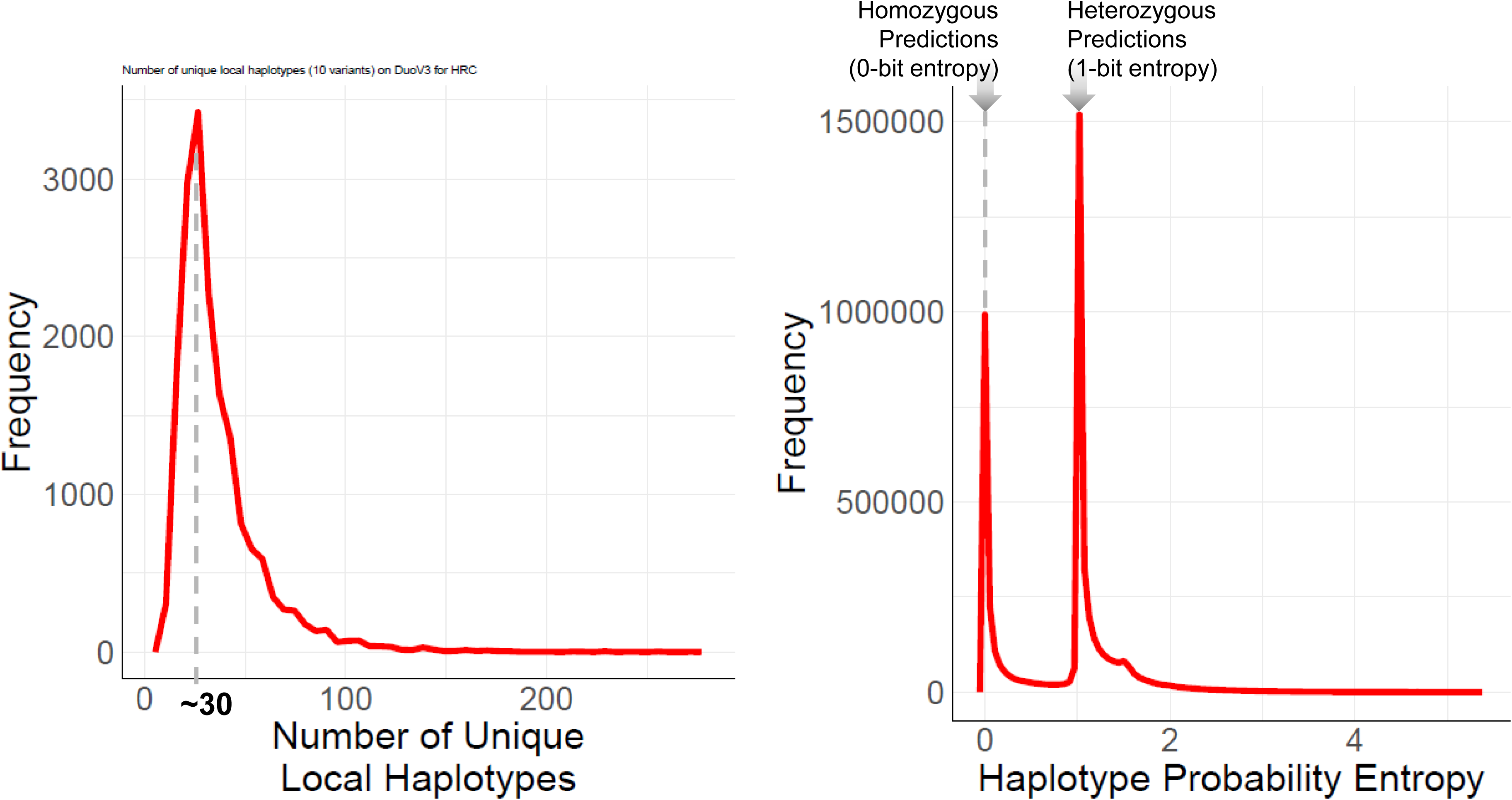

A simple interpretation of the local haplotypes within a block is that they represent a sparse embedding basis for the untyped variants in and around the locus of the local haplotype block. An important consequence of this is that the local haplotype of a subject can be used to “encode” and impute all variants in the vicinity of the local haplotype block.

For each block, we define the typed variants in the *n_vic_*-vicinity of the block as the haplotype classification features, which are the predictor features for the local haplotypes. By default, we use *n_vic_* = 100 with maximum stretch of 1 centiMorgan. The genetic distance cutoff enables focusing on regions with uniform recombination rates.

Next, the training data are generated for each untyped variant block. Since the reference panels vary by sample size (The 1000 Genome Project versus Haplotype Reference Consortium), it is necessary to augment the data and increase size. For each untyped variant block, the local haplotypes are sampled and paired randomly to generate the training data matrix. For each local haplotype, we store the indices for the unique local haplotypes (Fig. 1b), which serve as the target classes for training, which are encoded as the local haplotype probabilities corresponding to each haplotype. For each augmented pair of training haplotypes, the corresponding predictor-typed variant alleles are summed to generate predictor features. We store the unphased genotypes of the typed variants (i.e., coded as {0,1,2}) for the current block. For each block, we augment the training data size to 100,000 examples. The data are stored in binary formatted matrices for fast access. In addition to the training data, the unique local haplotype matrix is stored for each block.

For each block, a standard 3-layer MLP is trained. The first 3 layers are activated by rectified linear units (relu) and the last layer is designated to assign the haplotype probabilities (Fig. 1b). In the training phase, we used dropout layers (rate=0.25) to increase training robustness. Each MLP is trained for 10 epochs using ADAM optimizer and sparse cross-entropy loss. The models are converted to text-formatted matrices and stored for secure evaluation and processing. The haplotype matrices are stored for downstream analysis and are not immediately necessary for analysis.

At this point, it is worth emphasizing that the classification models are not trained to optimize genotype imputation accuracy. Thus, it may be expected that the trained models may not immediately provide high imputation accuracy. The models are trained to learn to discern between local haplotype classes, which give important intermediate information that SEmputica uses to develop further flexible downstream analysis tasks. Interestingly, our results show that these models provide much higher accuracy compared to the linear regression models while requiring same computational resources.

Genotype imputation is one of the most immediate applications of the haplotype classifier output models because all of the information necessary for imputation is available after model training. Given typed genotype matrix (e.g., VCF), SEmputica converts and gets the panel ready for imputation (e.g., importing, filling the missing genotypes). We first evaluate the trained models on the typed variants to obtain the probabilities for the local haplotypes. Given a genomic block, this step calculates, for each subject, a probability vector on the local haplotypes. The probability vector is projected on the haplotype matrix to obtain the final imputed genotypes (Fig. 1c,d). For each variant, *Dosage R^2^* (DR2) is calculated using the predicted dosages as a quality metric for the variants (Methods). SEmputica’s DR2 is used to filter out low quality variants. In addition to the conventional variants-level qualities, we perform subject-level filtering with low-quality local haplotype classifications using the entropy of the predicted local haplotype probabilities (Fig 2b).

#### Secure Evaluation of Haplotype Classifiers

The secure evaluation of the models can be efficiently performed by downloading the models locally to the querier’s site, which by definition protects client’s panel since it never leaves the site (Model to Data, M2D[26,33]). Since the models are generally very small in size, the clients can download the models and evaluate them locally with no data exchange with the server (Fig. 1c,d).

The simplicity of models and the flexibility of the imputation helps mitigate different types of new risks, e.g., *individual participation is revealed from variant locus only information*[32]. By splitting the imputation task into 2 steps, (1) haplotype classification and (2) final genotype prediction, SEmputica uses a simple form of *split-inference* (i.e., inference stage of split-learning frameworks[34,35]) where the last step of imputation is censored from clients while models are being shared. Further, releasing only the haplotype classification models opens new and flexible routes based mainly on *model obfuscation* to protect the reference panel in new ways.

For instance, the server can efficiently permute the haplotype classes for each subject (or buffer these) and release them to different clients. The models can be protected at the server side before being released: The server may choose not to release some of the haplotype classes by modifying/censoring the model parameters accordingly. Further, the server can introduce new haplotype states to obfuscate the real local haplotypes. Since local haplotype classes are not immediately linkable to other datasets by the client, these anonymized release mechanisms of model parameters help limit and manage leakage from the model parameters. The predicted haplotype probabilities can be used in downstream tasks (e.g., GWAS) since they are immediately interpretable in the context of genetics/biology.

When the client requests the actual imputed genotypes, the haplotype predictions are sent to the server after encrypting with CKKS scheme. The server performs the matrix multiplication of the haplotype probabilities with the haplotype matrices that are stored at the server side. This operation incurs only 1 inner product (similar to linear models[19]), i.e., SEmputica does not increase the complexity of the operations over the existing secure outsourcing models.

### Genotype Imputation Accuracy

We first compare the genotype imputation in three different settings on numerous array platforms.

#### Multi-Ancestral Testing Sample from The 1000 Genomes Project

We first use the 1000 Genomes Phase 1 release for testing imputation on a diverse panel. Among the 26 populations, we selected 20 subjects from each population and used a total of 520 subjects as the testing panel. The remaining 1984 subjects were used for training the haplotype classification models on chromosome 20. To assess the accuracy of different typed variant sets, we used seven different array platforms with varying coverages. We calculated genotype-level R2 as the accuracy metric to compare SEmputica’s imputed variants with our previous secure linear regression model (OLS) and BEAGLE as baseline models. To integrate the prediction quality with the method comparisons, we compared the variants that are deemed to be predicted accurately by all methods. We selected two cutoff’s for DR2 (0.5 and 0.8) and used the variants that are predicted above this cutoff by all methods.

Overall, we observed that SEmputica provides a major improvement in genotype-R2 when compared to OLS (Fig 3). The difference in genotype-R2 is more than 0.45 for especially the lower frequency variants. For all MAF categories, SEmputica’s imputations are always higher in accuracy than OLS, indicating that SEmputica’s haplotype-based models exhibit a major accuracy improvement compared to our previous secure regression models. When compared with BEAGLE, we observe that the variants in the MAF category 1-2% still exhibit lower accuracy, especially for the low coverage arrays. For the two lowest coverage arrays with less than 1 million typed variants, Global Screening Array (GSA) and UKBB-Axiom, genotype R2 difference between BEAGLE and SEmputica is around 0.12 (base) and 0.08 (tuned). On DuoV3b, Global Diversity Array (GDA), and Multi-Ethnic Global Analysis (MEGA) (more than 1.2 million typed variants), the genotype R2 of BEAGLE is 0.085 and 0.05 higher compared to base and tuned imputations of SEmputica, respectively. For the higher coverage arrays Omni2.5m and Omni5.4m, the BEAGLE still outperforms SEmputica’s imputations by 0.035 (base) and 0.025 (tuned) in genotype-level R2. When comparing common variants, we observed that the genotype R2 difference is less than 2% between BEAGLE and SEmputica.

**Fig 3:**
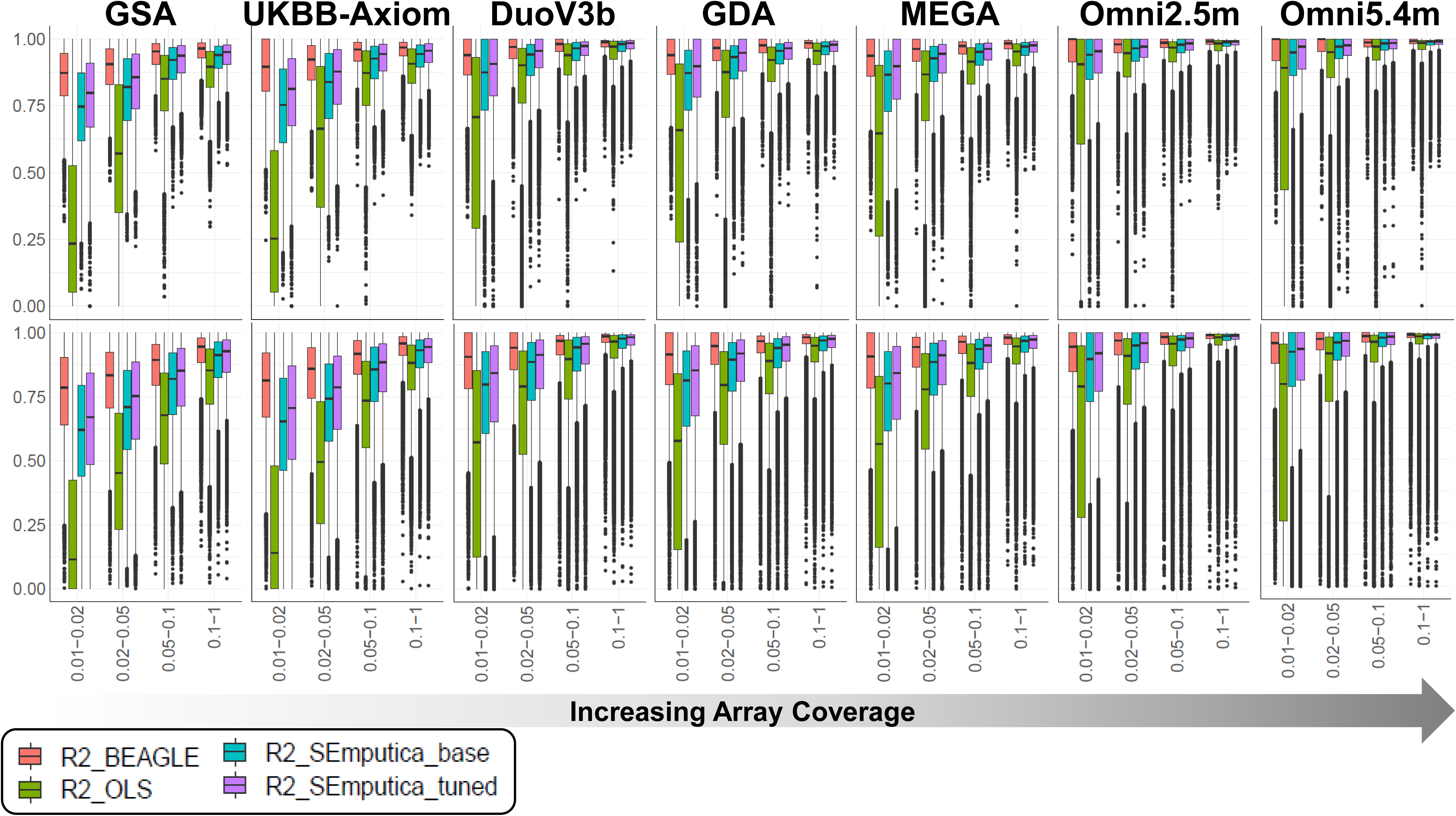
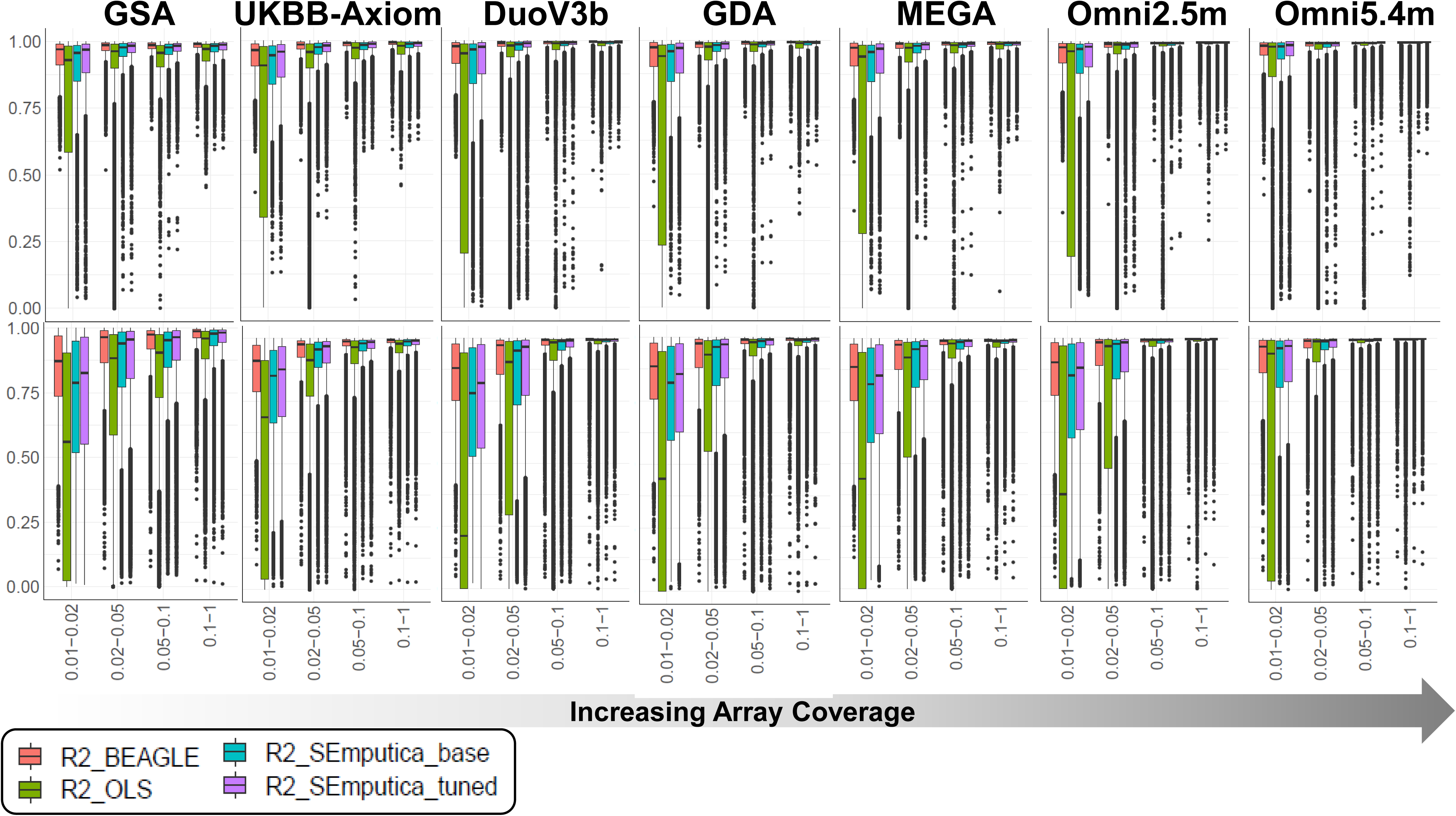
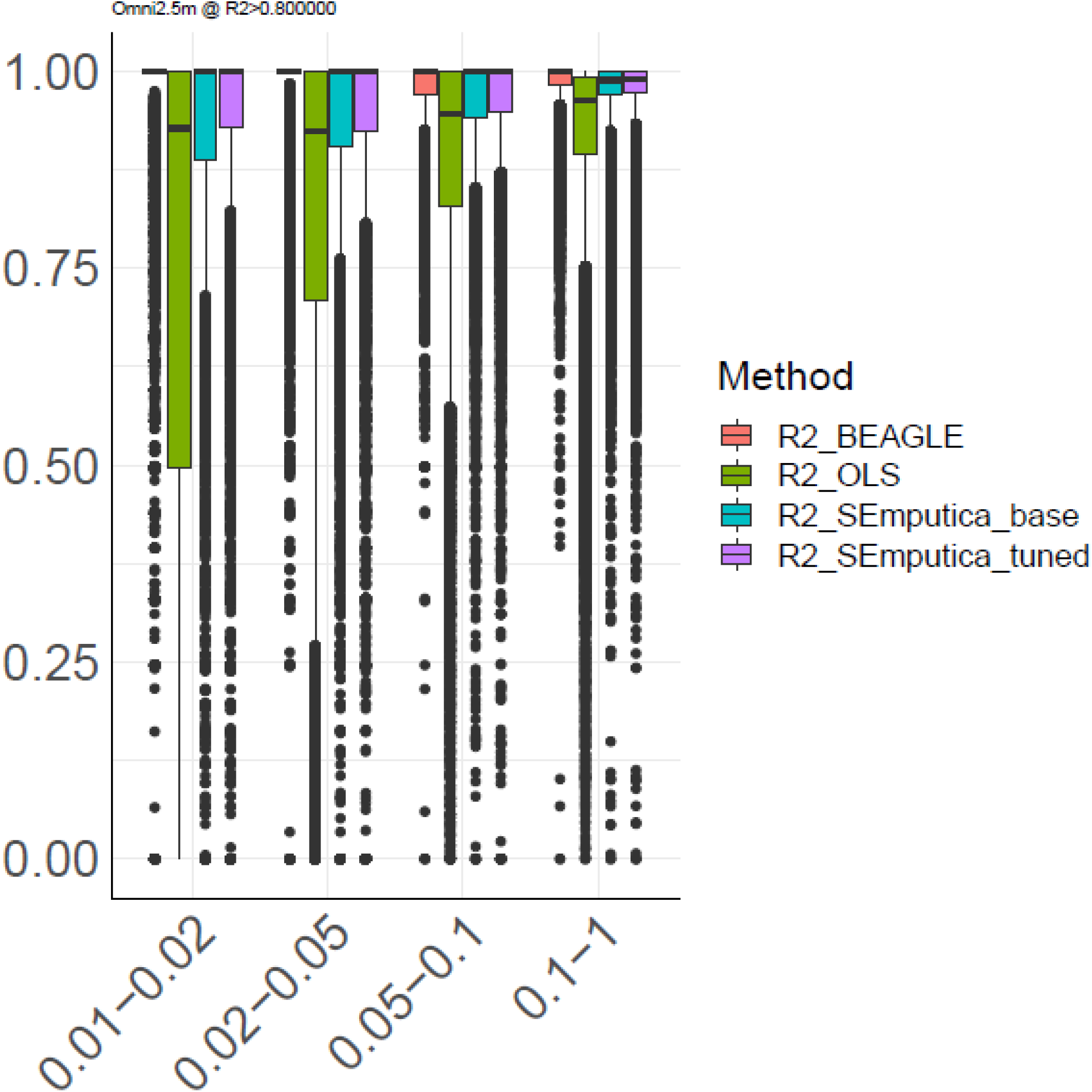

Overall, these results demonstrate that SEmputica’s local haplotype classifiers provide much more accurate genotype imputations than the linear models. However, multi-ancestral predictions using SEmputica can be challenging except for high-coverage arrays. It is important to emphasize that the low-frequency variant accuracies are generally not very robust for the small sample size we are using here.

#### European Testing Sample from Haplotype Reference Consortium Panel

To simplify the prediction scenario, we focused on the 24,576 European subjects in the Haplotype Reference Consortium dataset. We selected 5,000 random subjects to be used as the testing samples. The remaining 19,576 subjects were used to build imputation models on chromosome 20. We finally compared genotype-level R2 between the BEAGLE, OLS, and SEmputica’s base and tuned imputations (Fig. 3b).

The genotype level-R2 difference between BEAGLE and SEmputica is less than 5% across most MAF categories and for almost all array platforms, including the low-coverage array platforms. We still observe that the high coverage array platforms exhibit more even performance. For example, BEAGLE-vs-SEmputica exhibits less than 0.04 (base) genotype-R2 on Omni2.5m and around 0.021 (base) and 0.014 (tuned) on Omni5.4m for the lowest uncommon variants category. For common variants, the accuracy difference is below 2% for all array platforms. In comparison to OLS-based models, SEmputica’s imputations are higher with 0.20 in genotype-R2 for the lowest uncommon category that we tested.

#### Imputation of GTEx Panel

We finally used the 243 subjects genotyped on the Omni2.5m platform in GTEx Project and compared the imputed variants with the whole genome sequencing datasets. This case gives us a chance to evaluate whether prediction accuracy is robust with respect to the missingness that we observe on real array datasets. For this test, we used the models trained on the 1000 Genomes Project Panel. On more than 20,000 local haplotype blocks on chromosome 20, GTEx Omni2.5m array data contains around 2.5% missing genotypes. The accuracy different between SEmputica and BEAGLE’s imputations are generally similar as in our comparisons: In the lowest uncommon variant category, BEAGLE slightly outperforms SEmputica’s base imputations with 0.035 R2 difference and tuned imputations by 0.03 (Fig. 3c).

### Resource Requirements

The model database uses 1.2 Gigabytes of disk space, which includes the text-formatted model parameters (extrapolating to the whole genome is approximately 60Gb).

### Variant and Local Haplotype-based Association Testing

We next asked if the SEmputica’s haplotype classification and genotype imputation can be used in downstream analysis, specifically in a genome-wide association study (GWAS). We simulated continuous phenotypes on European testing panel from HRC using causal variants from within imputed variants on Omni5.4m panel. We focused on two scenarios with phenotype heritability *h*^2^ = {0.5, 0.8}. After simulating phenotypes, we used the variants imputed by BEAGLE, SEmputica-base, SEmputica-tuned and estimate the summary statistics using plink2[36] using 10 PCs to correct for account for confounding by population stratification. This process was repeated 100 times to test concordance between association testing statistics. Overall, the association testing p-values assigned for genotypes imputed by SEmputica against those imputed by BEAGLE, for the known variants have high concordance (Pearson R^2^>0.98, Fig. 4a), indicating that the summary statistics calculated using the SEmputica imputed variants are highly concordant with those estimated using BEAGLE imputed variants and the original sequencing-based variants.

**Fig 4:**
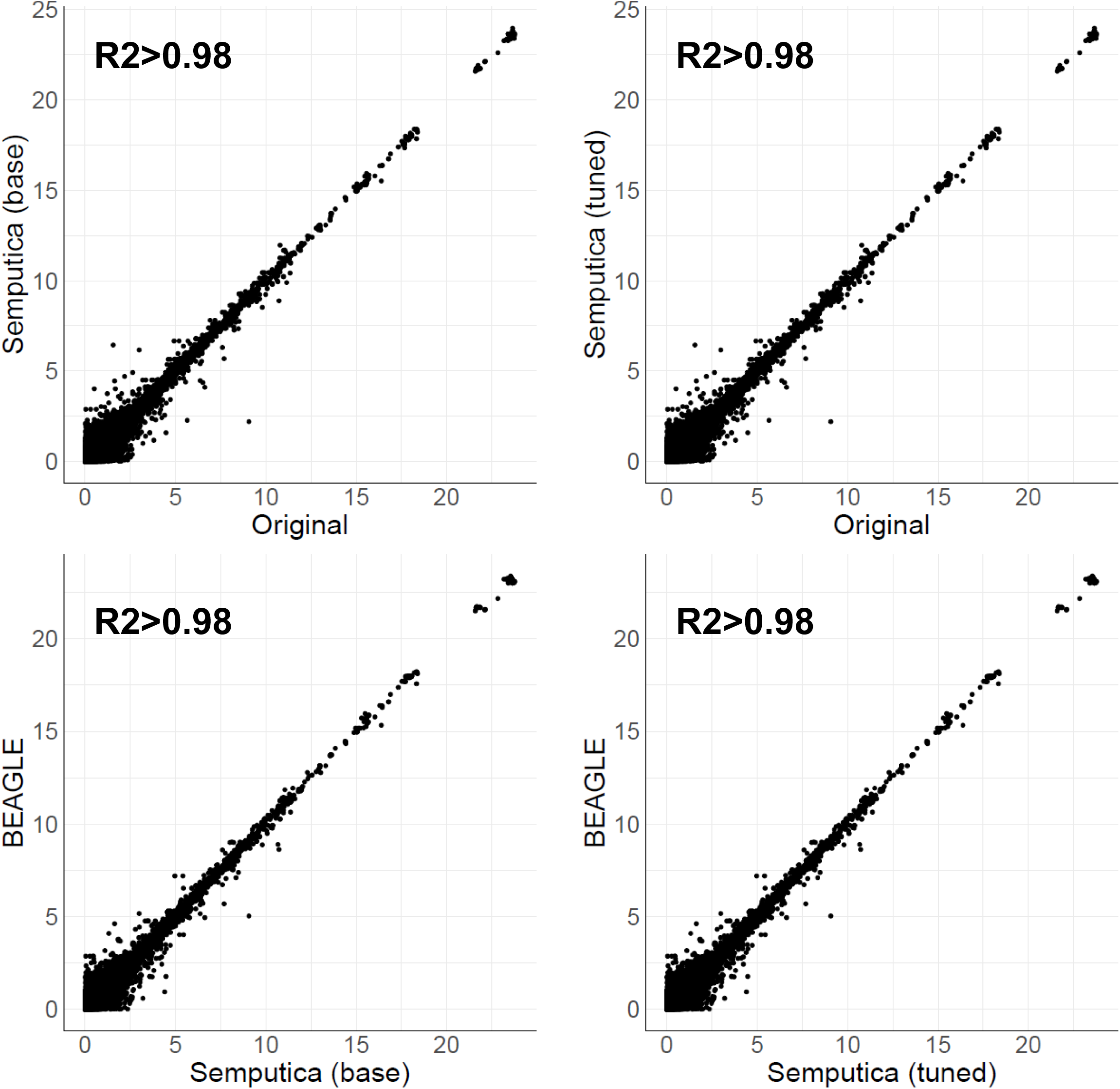
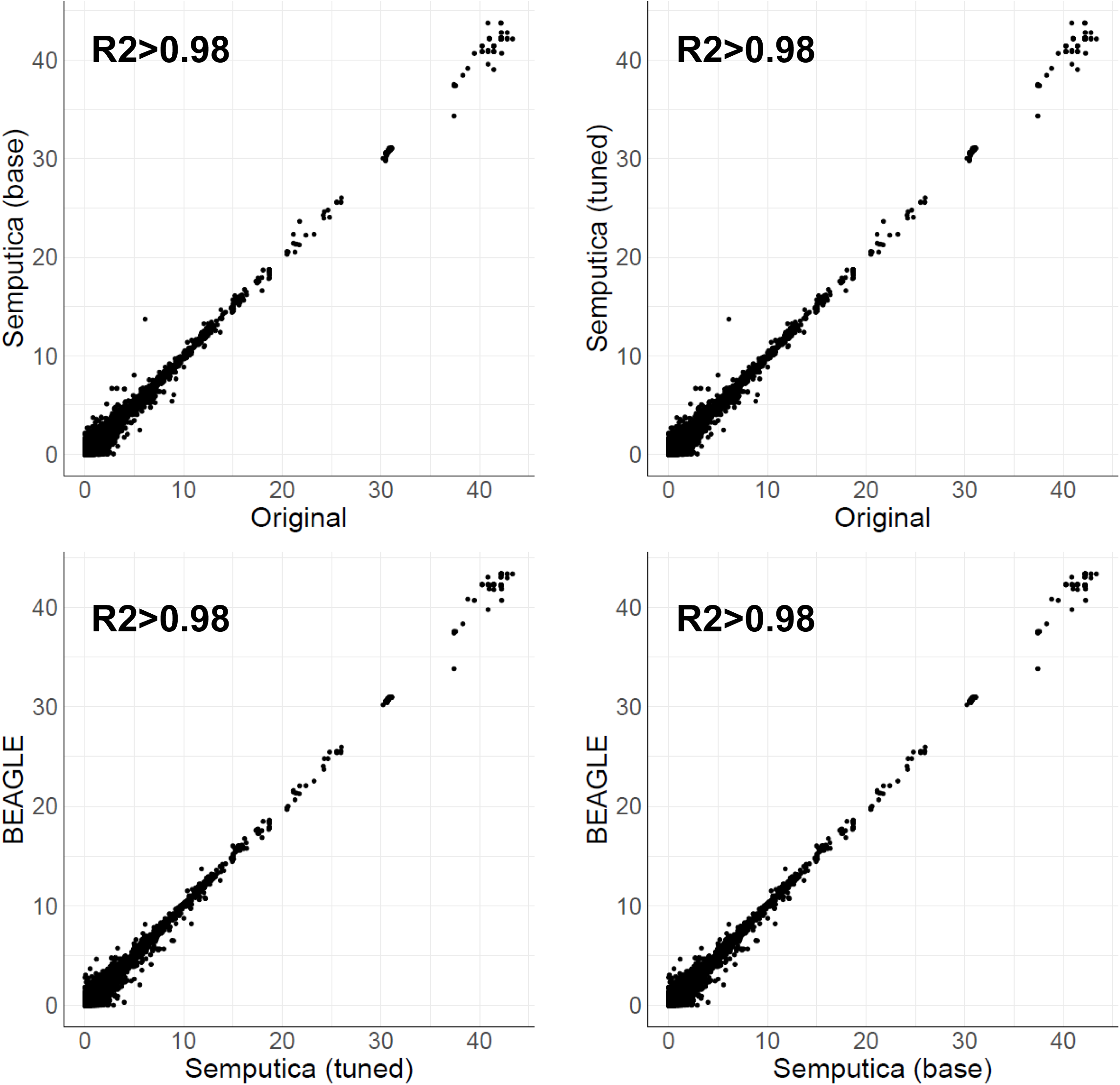
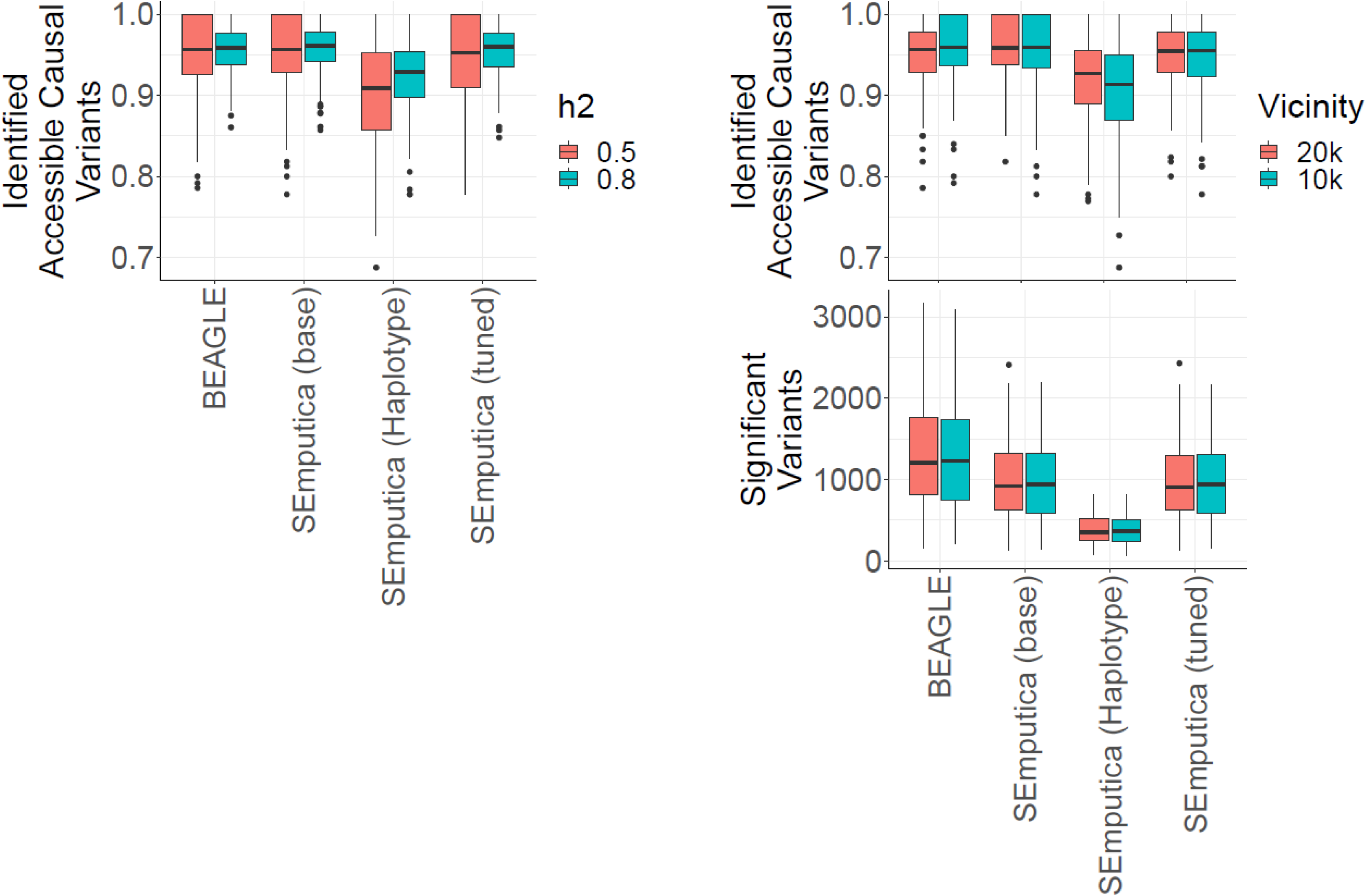

We next asked if local haplotype probabilities can also be used for performing GWAS and downstream collaborations. This is advantageous for privacy preservation since the client can calculate haplotype probabilities locally without any interaction with a server. We propose using haplotype probability matrix to perform GWAS directly in a linear model (Fig. 1c). Unlike variant-based association testing, the haplotype-based tests[37] model the contribution of each local haplotype (with probability between 0 and 1) with a linear regression coefficient. After model fitting, we test the significance of the coefficients assigned to the haplotypes. Haplotype-based testing provides a flexible route for privacy protection and collaboration between sites since we perform associations using local haplotypes (and not the actual variants).

We ran the local-haplotype-based association tests using haplotype probabilities predicted by SEmputica among approximately 20,000 blocks on Chromosome 20 using the simulated phenotypes used in previous setting. We next calculated the number of variants that are in the vicinity of the *accessible causal variants*. By accessible causal variant, we mean the causal variants whose plink2-assigned association p-value is smaller than 10^-8^ and are imputed by BEAGLE dosage R2 greater than

0.8. Among the accessible causal variants, we calculated the fraction of variants and blocks that are identified as significant in association testing (p-value < 10^-8^) (Fig. 4c). As expected, SEmputica imputed variants can identify a similar fraction of the accessible causal variants as BEAGLE. In comparison, local haplotype-based tests of SEmputica identifies slightly lower fraction of variants (3% of accessible variants are missed by haplotype-based tests for h^2^=0.8 at 20,000 base pair vicinity of the accessible variant). We also observed that the haplotype-based tests provide a much smaller number of associated regions compared to the variant-based testing counterparts.

## Discussion

We presented SEmputica, which comprises of haplotype classification models and downstream analysis tools. We have three main objectives in this study: First, we present SEmputica’s haplotype classification-based framework as a basic tool for building HE-capable haplotype analysis, imputation, and downstream analysis methods. Compared to the previous secure models that relied on the linear models that were only capable of imputing one untyped variant at a time, SEmputica extends the frontier of secure genetic analysis by integrating haplotype-based analysis, imputation, and numerous downstream tasks (e.g. GWAS) into one framework.

SEmputica’s classification models can be approximated by low-depth circuits and are amenable for HE-conversion (i.e., HE-portable). Thanks to the new developments in the HE literature, e.g., HE-capable secure neural network evaluations[38], we foresee that evaluations of such methods will be routinely possible on the commodity low-cost hardware with no limitation on the platform. When the clients evaluate the models locally on-site, the complexity of secure imputation using HE-based primitives is the same as running linear inner products (one matrix multiplication per block), which can run at deployment scale as we have demonstrated in our previous study.

SEmputica’s architecture is compartmentalized to simplify privacy protection: For example, the haplotype classification, imputation, and association testing modules are split so that computationally demanding steps, i.e., haplotype classification models, can be run at the client locally with no data exchange with a server, similar to the “model-to-data” philosophy for outsourcing computations. Although deep-learning-based methods have been proposed with the same privacy-protection philosophy, they require prohibitively large computational resources to train, creating almost a unique dependence on specific hardware platforms (e.g., NVIDIA GPUs). Moreover the deep learning models may be extremely large in size, which may limit their transfer rates due to network bandwidth limitations. SEmputica’s haplotype classification models are orders of magnitude simpler and can be trained on CPUs (although we do require tensorflow libraries) and evaluated on CPUs (with no reliance on tensorflow). Further, these models do not provide means to protect ultra-rare variant loci in the reference panels[32]. On the other hand, SEmputica mitigates this risk by sharing the models for haplotype classification, which does not reveal the actual loci of the untyped variants in the panel. These predictions reveal only blocks, whose positions do not have to be revealed to the client. This way, the reference panels add a unique layer of protection between the client and themselves. In case the client requests actual genotype imputations, the local haplotype probabilities are encrypted and sent to the server, which can decide which untyped variants can be sent. This way, the server now possesses the property of “*deferred release of variant loci*” strategy. We believe that this property can reveal new opportunities for releasing haplotype-level data without the need to release actual variant loci or genotypes by releasing the haplotype classification models. For example, the imputation servers can only serve haplotype classification models by limiting large loci (e.g., HLA, APOE). While sharing the models with a client, the server can choose the limit of the haplotypes that are shared by simply aggregating them in the final layer of the MLP model parameters. The server can also randomly shuffle the local haplotype classes that are released to a client to anonymize them among different clients. Since local haplotypes are highly reference panel specific, it is unlikely to perform reliable linking attacks by cross-referencing local haplotype-based results even if the haplotype probabilities are accidentally released to unauthorized entities.

In essence, the reference panel haplotypes can be considered as interpretable embeddings of the actual variant genotypes. This is because variants do not exist independently of each other and manifest on the diploid haplotypes. SEmputica aims to use this biological property and uses reference panel-specific local haplotypes as units of sharing genetic information. Local haplotype probabilities can be used for further downstream analysis in a haplotype-based GWAS analysis, which have also been shown to be more sensitive than variant-based GWAS methods.

As we discussed above, SEmputica haplotype classification models are designed with a focus on simplicity and compartmentalization in mind to enable privacy and HE-portability. Our results show that this reliance on simplicity may penalize imputed genotypes in terms of accuracy when compared to BEAGLE. However, we found that SEmputica’s imputations exhibit a major improvement compared to the linear models, which were the only viable models for secure genotype imputation. Further, although the accuracy difference in multi-ancestral panel tests may be high, our results show that the accuracy of reliable variant imputation on European populations is highly comparable to BEAGLE’s imputations. Considering that most cross-ancestry analyses are performed by an initial focus on the ancestry-specific analyses followed by an integrative analysis (e.g., meta-analysis), SEmputica’s future development will largely focus on building more accurate ancestry-specific models followed by downstream tools that aim at integrating results from different ancestries[39]. Considering that the rare variants tend to be highly population-specific (e.g., due to bottleneck events), it is reasonable to expect that the haplotypes that harbor these variants will be more accurately modeled using ancestry-specific models.

At this point, we would like to reiterate that SEmputica’s models are trained to maximize haplotype prediction accuracy, and genotype imputation accuracy is not optimized while training the models. This is a deliberate decision as we view SEmputica’s local haplotype classification-based framework aims at enabling a privacy-aware haplotype-level analysis. Within this framework, genotype imputation stands as a specific application. As with any predictive method, genotype imputation and related methods and data should be treated carefully. For instance, the imputation accuracy relies heavily on the ancestral composition of the reference panel’s and testing panel’s ancestral compositions, the selection of the genotyping array platform[40], and parameters of imputation software. While imputed genotypes are used in downstream tasks such as GWAS and meta-analysis, it is highly advisable to select stringent thresholds on the imputed variants, especially for the low-frequency and rare variants. We believe that it is very important to realize that variant imputations are computationally derived estimates of genetic information and should be treated as such. Usage of high-density arrays is highly advisable for analysis that will focus on rare variants.

## Methods

We describe the details of methods used to train and test SEmputica’s models.

### Local Haplotype Classification Model Training Data Augmentation

Given a phased reference panel, SEmputica first re-samples the phased panel using ProxyTyper to increase haplotypic diversity. For 1000 Genomes Project data, we resampled data to 10,000 subjects. This resampling step uses all of the variants. For Haplotype Reference Consortium panel, the resampling step was not used due to the large size of the dataset.

Next, variants are filtered with respect to minor allele frequency (MAF) cutoff. For high coverage arrays, we filtered variants with MAF lower than 0.01. This is applied to both the typed and untyped variants. For typed variants, MAF filtering removes the predictive features (i.e., tag variants) that show very small variance. While HMM-based methods can seamlessly make use these almost non-informative typed variants (since they do not operate on small local windows), regression-based methods make better use of features that have higher variance.

For the untyped variants, we use the same MAF filtering cutoff. After this filtering, we use the reference panel blocks that are used to define the local haplotypes at each locus. By removing the rare untyped variants on the reference panel (MAF<0.01), we cut down the number of unique local haplotypes in each haplotype block.

Next, the untyped variants are used to define the local haplotype blocks. We use 10 consecutive variant blocks for The 1000 Genomes Project and the Haplotype Reference Consortium panels.

### Local Haplotype Classifier Models and Training

For each block, the unique local haplotypes of the untyped variant in the block are computed. At each block, we randomly sample haplotype pairs. We add the selected pair of haplotypes for the typed variant (input features) that are within *n_vic_* vicinity of the untyped variant block (*n_vic_=100* by default). We store the pair of indices for the the sampled haplotype pairs among the unique local haplotypes and the typed variant unphased genotypes as training examples for the classification models.

The classifier models are structured as 3-layer dense networks (MLPs) and are trained using keras with tensorflow2 backend. Rectified linear units were used as activation functions and dropout layers were added with rate=0.25 to robustify trainings. The output from MLPs represent the unique haplotype probabilities for the input typed variants.

### Dosage R^2^ Estimation for Regression-based Models

For each variant, we calculate the dosage R^2^ (DR2) as the fraction of the variance in dosage and imputed genotype variances. After assignment of DR2, we recommend filtering DR2>0.8 to flag these variants as low-quality variants.

### Local Haplotype-Based Association Testing and Meta-Analysis

For each block, we use the local haplotypes to quantify the delineation of phenotypes on the Haplotype-based GWAS is calculated using the per haplotype probability output from MLP. For each subject with phenotype (*y*), we fit the linear models to decompose phenotypes on block-*i*:

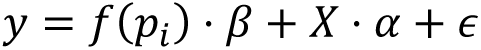

where *y* denotes the phenotype (continuous/binary) vector, *X* denotes the covariates matrix (rows: Subjects, columns: covariate features) and *ε*∼*N*(0, σ*_e_*) denotes the independent phenotype component, *p_i_* denotes the haplotype probability matrix and *β* denotes the coefficients for the haplotype probabilities on the phenotype. *f*(*p_i_*) denotes a suite of linear/non-linear operators to summarize when there are large number of haplotypes (columns) in the block.

SEmputica reports standard errors, effect sizes, and p-values that can be used to perform further downstream analysis such as secure meta-analysis of different studies in collaborations.

